# Auditory cortex activation is modulated nonlinearly by stimulation duration: A functional near-infrared spectroscopy (fNIRS) study

**DOI:** 10.1101/2021.08.02.454752

**Authors:** Yi Fan Zhang, Anne Lasfargue, Isabelle Berry

## Abstract

Functional near-infrared spectroscopy (NIRS) is an increasingly popular method in hearing research. However, few studies have considered efficient stimulation parameters for fNIRS auditory experimental design. The objectives of our study are (1) to characterize the auditory hemodynamic responses to trains of white noise with increasing stimulation durations (8s, 10s, 15s, 20s) in terms of amplitude and response linearity; (2) to identify the most-efficient stimulation duration using fNIRS; and (3) to generalize results to more ecological environmental stimuli. We found that cortical activity is augmented following the increments in stimulation durations and reaches a plateau after about 15s of stimulation. The linearity analysis showed that this augmentation due to stimulation duration is not linear in the auditory cortex, the non-linearity being more pronounced for longer durations (15s and 20s). The 15s block duration that we propose as optimal precludes signal saturation, is associated with a high response amplitude and a relatively short total experimental duration. Moreover, the 15s duration remains optimal independently of the nature of presented sounds. The sum of these findings suggests that 15s stimulation duration used in the appropriate experimental setup allows researchers to acquire optimal fNIRS signal quality.

## Introduction

Functional near-infrared spectroscopy (fNIRS) is an essential emerging tool for functional monitoring and imaging of brain hemodynamics used to study human brain function (Boas et al., 2004). As brain activity induces a local increase in oxygen consumption, fNIRS is able to spectroscopically characterize the concentration of oxygenated (HbO), deoxygenated (HbR), and total hemoglobin (HbT) within the brain exposed to near-infrared light in a range of wavelengths between 650 and 860 nm, in which photons can penetrate the intact scalp to illuminate the cerebral cortex. Using the modified Beer-Lambert law, we can calculate changes in the amount of diffuse light reaching the detector, which corresponds to changes in HbO and HbR concentrations in the tissue (Boas et al., 2004). As in functional magnetic resonance imaging (fMRI), the changes in the concentration of HbO and HbR are considered an indirect measure of brain activation, reflecting the relation between neuronal activation and metabolic demand based on neurovascular coupling (Steinbrink et al., 2006). Because of its safety (noninvasive and nonionizing nature), portability, ease of application, and moderate level of resistance to movement artifacts, the technique is particularly suitable for vulnerable populations including infants, children, and patients for whom other imaging methods are limited (Ferrari and Quaresima, 2012). Unlike fMRI, the operation of fNIRS is completely silent and unaffected by magnetic interference from cochlear implants. Most importantly, fNIRS can circumvent the limitations of acoustic scanner noise (Peelle, 2014), thereby enabling its extensive research applications in the field of auditory neuroscience (Fava et al., 2014; Pollonini et al., 2014; Remijn et al., 2017; Basura et al., 2018; Hiwa et al., 2018; Anderson et al., 2019). Furthermore, it offers unprecedented opportunities to acquire high-quality data for hemodynamic responses to auditory stimuli.

However, for fNIRS to become an effective tool for studying auditory functions, it is critical to take into consideration the significant variability in experimental designs and analysis techniques across fNIRS studies, as these can make the interpretation and replication of studies difficult (Herold et al., 2018; Issard and Gervain, 2018; Pinti et al., 2019). To develop more easily replicable and more efficient experimental designs, researchers should consider a range of factors in their stimulus presentation protocol, which affects the resulting hemodynamic response (HDR). That the characteristics of the evolving HDR vary as a function of stimulus properties and experimental parameters has been well-established. A good knowledge of the interrelationship between stimulus parameters and the functional imaging signal response is critical for designing the experimental protocol and interpreting cortical activation. Initially, one should carefully consider auditory stimulus properties, which include intensity, frequency modulation, single stimulus duration, and spatial cues (Bauernfeind et al., 2018; Weder et al., 2018, 2020; Zhang et al., 2018). Then, stimulation parameters should be optimally chosen, including stimulus presentation rate (Binder et al., 1994), interstimulus intervals, and block stimulation durations (Robson et al., 1998; Hu et al., 2010). Stimulation properties should be considered in the context of the sampling procedure of HDR (Miezin et al., 2000). However, to date, there is no consensus regarding the most-efficient stimulation parameters for fNIRS experimental design.

In some studies, both fMRI and fNIRS demonstrated a linear relationship between the stimulus parameters and the HDR (Boynton et al., 1996; Wobst et al., 2001). With the assumption of linearity, the response to multiple stimuli might be predictable by the superposition of the responses to each individual stimulus (Robson et al., 1998). However, the results across other studies are contradictory. Among the stimulus parameters, the effects of the stimulus duration on HDR vary widely in different sensory modalities (Soltysik et al., 2004). For example, Robson et al. reported that the HDR in the auditory cortex to trains of tones is approximately linear for trains of 6s and longer but nonlinear for less than 6s (Robson et al., 1998). Similarly, Soltysik et al. observed nonlinearity in the primary auditory cortex for stimuli of less than 10s and a more linear increase for longer stimuli. Conversely, Glover demonstrated that the auditory response is linear for short-duration auditory stimuli but highly nonlinear for longer stimuli (Glover, 1999). The authors showed a higher degree of nonlinearity in the blood-oxygen-level-dependent (BOLD) response changes in the visual cortex for shorter-duration stimuli (Vazquez and Noll, 1998; Liu and Gao, 2000; Pfeuffer et al., 2003). A nonlinearity response is also shown in the human somatosensory cortex (Nangini et al., 2002).

To the best of our knowledge, fMRI was used as a tool in almost all the studies that investigated the relationship between stimulation duration and HDR, and no result using fNIRS has been reported in relation to this topic in auditory research. Given the critical advantages provided by fNIRS in the field of auditory neuroscience compared with fMRI, our research aims to study the effects of auditory block stimulation duration on HDR as measured by fNIRS. In addition, we assess the linearity or nonlinearity of the HDRs induced by different stimulation durations in the primary auditory cortex. This analysis aims to improve the design of fNIRS auditory experiments when task duration varies (Soltysik et al., 2004) and to increase the understanding of brain adaptation peculiarities (Krekelberg et al., 2006).

Furthermore, although the effects of stimulus duration have been extensively reported in previous studies using fMRI, no standardized stimulation duration has yet been proposed. As the signal-to-noise ratio (SNR) is lower than in fMRI (Cui et al., 2011), block designs are more widespread in fNIRS studies for maximizing the signal-to-noise ratio and increasing the detection power of activation (Amaro and Barker, 2006; Shan et al., 2014). According to dominant application areas of fNIRS in infants and children, as well as in neuropsychiatric patients (Boas et al., 2014), the longer experimental duration can lead to uncooperative and fidgety test subjects. In practice, experimental duration is a crucial factor to be considered. Thus, the second objective of our study is to identify the best auditory stimulation duration in the block design that produces the most-efficient activation with maximal amplitude in a minimal duration as the most optimal choice for future applications.

In our study, we first localized the HDR to auditory stimuli. For this purpose, we chose a given number of auditory stimuli and distributed them among four conditions of stimulation duration that are most commonly employed in task-presentation protocols for auditory studies. Next, we compared the amplitude of the induced HDR and the total experimental duration for each condition and then suggested the most-efficient stimulus duration that could be generalized to other types of auditory stimuli. Moreover, we investigated in further depth the relationship between HDRs and different stimulation durations through the assessment of linear additive effects. We used a variety of natural sounds and white noise in order to eliminate the assumption that results depend on stimulus type.

## Materials and methods

### Subjects

A total of 17 adult hearing subjects (9 males and 8 females) ages 27 ± 5 years (mean ± SD) participated in the experiment. All participants completed a self-assessment questionnaire; none had hearing disabilities; none had neurological disorders; all were right-handed; and all had normal vision or vision corrected-to-normal. Participants were divided into two experimental groups: 9 subjects for the first experiment where the stimulation duration varied (8s, 10s, 15s, 20s) and 8 for the second experiment where we contrasted two types of auditory stimulation (white noise/natural sounds).

All participants gave their informed written consent to taking part in the study and for the publication of the results in an online open-access database. The study was approved by the local ethics committee at Paul Sabatier University. The subjects were not paid for their participation.

### Experimental set and stimuli

The experiment was conducted in a sound-proof booth with a 27-inch screen and luminosity maintained at an attenuated stable level. The display was placed in front of the participant at a distance of 80 cm. The audio stimuli were presented at a sound pressure level of approximately 75 dB through anti-noise headphones (SONY) connected to an external sound card (MOTU). The experimental installation was surrounded by gray curtains.

Each participant performed a passive listening task. The presented stimuli were either white noise or vocal and nonvocal sounds according to different experimental conditions. All stimuli were presented binaurally and lasted 500 ms. In addition, the intensity was equalized according to the root mean square level. The vocal and nonvocal stimuli used in this study originated from the database exploited in previously published studies (Belin et al., 2000; Massida et al., 2011; Vannson et al., 2020). They comprised speech stimuli and nonspeech vocal stimuli (e.g., laughs, coughs) and environmental sounds, including sound alarms, car horns, streaming water, animal vocalizations, etc.

### Experimental design

The experiment was divided into two parts. In the first part, we varied the block stimulation duration, but the total number of presented stimuli remained nearly unchanged (the number of blocks * the number of stimuli per block = total number of presented stimuli) and the presentation rate per second was the same across conditions. There were four stimulation conditions according to the duration of stimulation, including 8s, 10s, 15s, and 20s. Thus, a block design was implemented for each condition. Details of stimulation proprieties are indicated in Table 1. The presented stimuli were white noise through all four conditions. Each subject experienced, in randomized order, the above-described conditions so that each condition was presented one time with a 1-minute pause between the conditions.

**Table 1.**
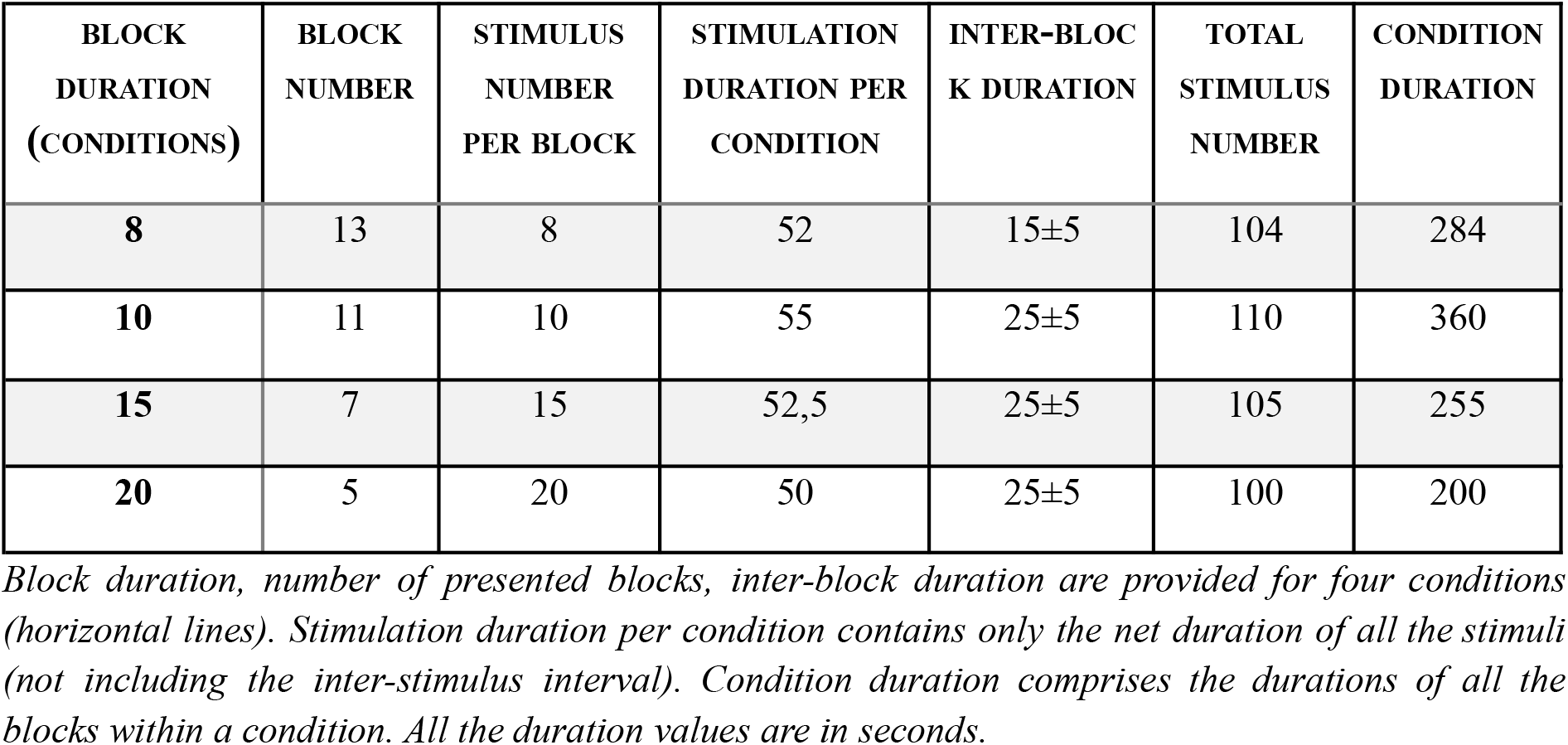
Stimulation properties of experimental conditions.

The experimental design is shown in Fig. 1. At the start of each condition, 60 seconds of silence was presented as a baseline. After that, participants were presented with the number of blocks of white noise corresponding to each condition. We introduced a jittered interblock interval in the range of 15 ± 5s or 25 ± 5s. Both of the above time windows were chosen to be of sufficient duration for the descent of the BOLD signal to baseline (Malonek and Grinvald, 1996). The overall experiment lasted approximately 25 minutes. During that time, participants were instructed to look at the fixation cross. In the second part of the experiment, we contrasted two types of stimuli: white noise and vocal and nonvocal sounds, applying the same stimulation parameters. There were two conditions: the white-noise condition and the vocal/nonvocal sounds condition. Thus, eight other subjects performed the two conditions in random order.

**Fig. 1.**
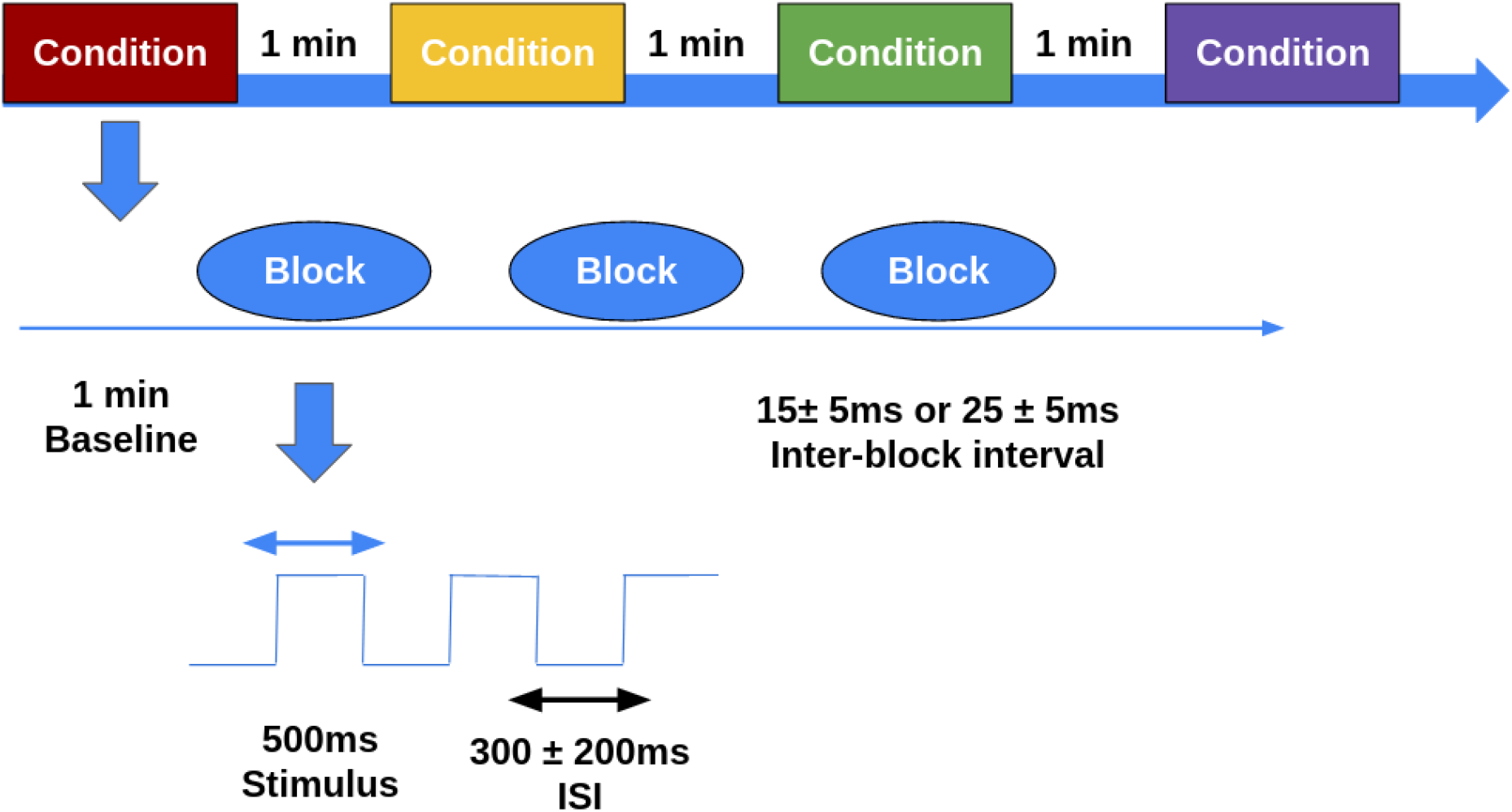
Experimental design. Three levels of experimental design, from top to bottom, the experimental level, condition level, and block level. Squares represent four experimental conditions and oval forms represent blocks of presented auditory stimuli (white noise) within the condition. The scheme below oval forms represents the time course of a block for a series of stimuli presentations with a variable inter-stimulus interval of 300±200ms, indicated by the black double arrow.

Before the above-described experiment, all participants were instructed to follow three simple directions: (1) listen carefully to the auditory stimuli; (2) remain still, keeping your arms on the armrests of the chair during all experiments; and (3) avoid the slightest head movements during the auditory stimulus presentation.

### Data acquisition

The hemodynamic change in subjects’ brains in response to the auditory stimuli was recorded using an NIRS device (NIRx NIRScout 816, NIRx Medizintechnik GmbH, Berlin, Germany) with a sampling frequency of 8 Hz. The instrument generated two different wavelengths (760 and 850 nm) and measured the time course of changes in oxyhemoglobin (oxy-Hb), deoxyhemoglobin (deoxy-Hb), and total hemoglobin, using the modified Beer-Lambert law. The probe set contained two 3 × 2 arrays (four sources and four detectors per hemisphere), resulting in a total of 20 measurement channels (Fig. 2a and 2b).

**Fig. 2.**
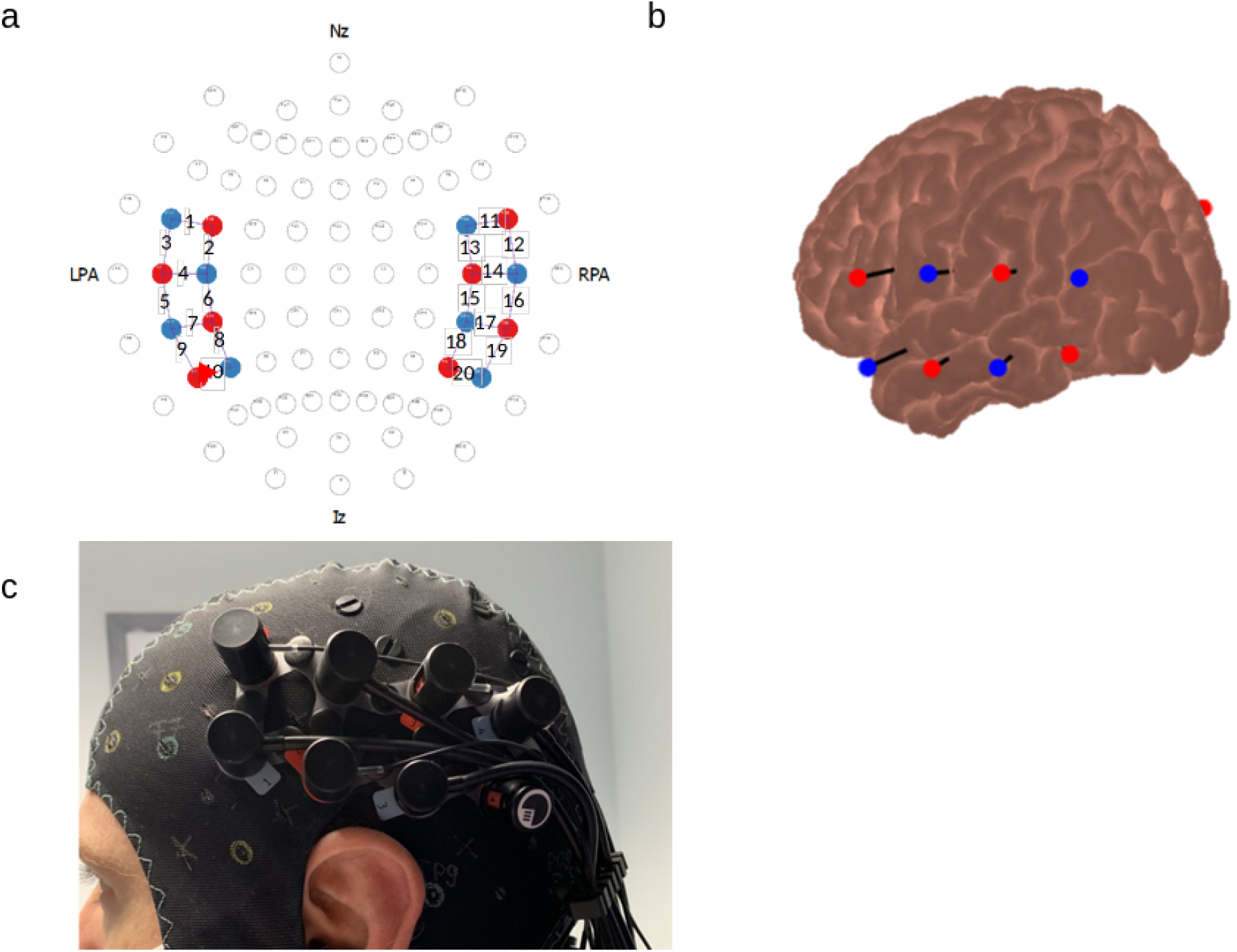
Schematic description of NIRx NIRScout 816 NIRS machine. **(a)** Probe montage overlaid on the scheme of the 10–20 system. Blue circles indicate detectors; red circles indicate sources; and numbered lines correspond to measurement channels. LPA and RPA: left and right preauricular. Nz: nasion; Iz: inion. **(b)** Projection of probe positions on the MNI brain surface. **(c)** Photograph of optode array placement on the standard NIRSx NIRS cap used for this study.

The pairs of source-detectors were mounted in a standard NIRX NIRS cap with a fixed distance of 3 cm. The standard NIRX NIRS cap had predefined EEG layout positions (Fig. 2c) in accordance with surface anatomical landmarks and following the international EEG 10–20 system (Homan et al., 1987). Placing the cap on the subject’s head was guided by the experimenter from a set of fiducial points (nasion, inion and left and right preauricular anatomical landmarks). Afterward, the experimenter ensured that the optodes were in contact with the participant’s scalp. To ensure test-retest reliability of fNIRS assessment between experiments, the experimenter confirmed that the cap was set up as consistently as possible across experiments and participants. The size of the standard NIRS cap was adjusted according to the head circumference of each subject for a valid intersubject comparison of cortical areas underlying the measurement channels.

The optode set (Fig. 2a–2c) was positioned on the head over the temporal and frontal cortex regions of both hemispheres to provide sufficient coverage of superior temporal regions, where different types of the auditory cortices are primarily located. To confirm that the probes were sensitive to the relevant brain areas, we used the Monte Carlo photon transport software tMCimg via AtlasViewer to estimate the probabilistic path of photon migration through the cortex for the sensitivity profile of the optodes (Boas et al., 2002; Aasted et al., 2015). Fig. 3 depicts the regions to which probes were sensitive, covering the middle and superior temporal regions, thereby confirming the validity of our probe set.

**Fig. 3.**
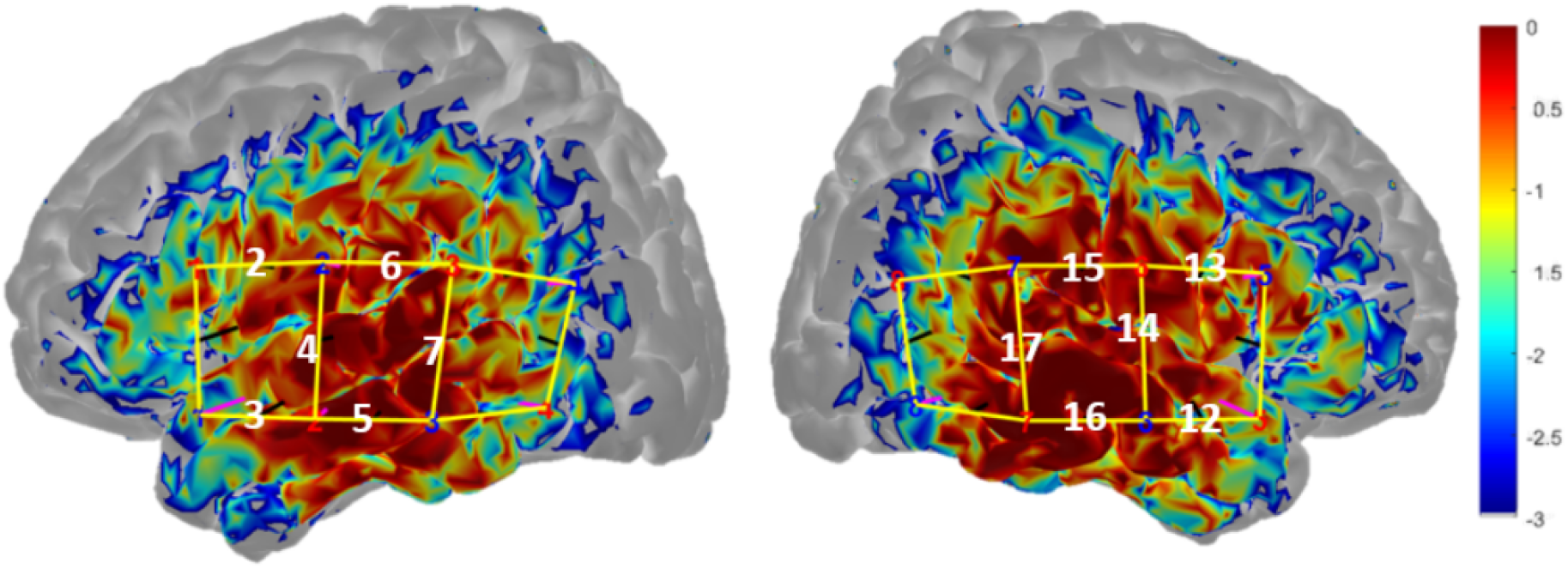
Results of Monte Carlo-simulation for the optode configuration used in our study. The color bar indicates the spatial sensitivity of fNIRS. It is expressed in mm^-1^ and values range from 0.001 to 1 in log10 units : -3 to 0. The measurement channels 2, 3, 4, 5, 6, 7 for the left hemisphere and measurement channels 12, 13, 14, 15, 16, 17 for the right hemisphere presented a good sensitivity to the underlying cortical areas.

For the subsequent processing and statistical data analysis, only the HbO values were utilized and illustrated in all tables and figures. The choice of HbO is based on results from previous studies, which demonstrated more-significant or more-robust effects with HbO than with HbR (Huppert et al., 2006; Ye et al., 2009; Gagnon et al., 2012; Issa et al., 2016) and higher correlation with the fMRI-BOLD response than HbR (Strangman et al., 2002). Therefore, the choice of HbO as an investigation parameter of the hemodynamic response allows us to better compare the results with previous functional neuroimaging studies, especially fMRI.

### Processing data analyses

The fNIRS data were processed in MATLAB (MathWorks, Natick, MA) using functions provided in the HOMER2 package (Huppert et al., 2009). In the first step, signals were converted from light-intensity values to optical-density levels using the hmrIntensity2OD function in HOMER2. Afterward, principal component analysis (nSV=0.8) (Zhang et al., 2005; Wilcox et al., 2008; Virtanen et al., 2009; Wilcox et al., 2010) was applied to the fNIRS data using the enPCAFilter function in order to separate the component of the HDR from undesirable sources and to improve the contrast-to-noise ratio (Zhang et al., 2005). Motion artifacts were suppressed using the hmrMotionArtifactbyChannel function (tMotion = 0.5, tMask = 2.0, STDEVthresh = 20, and AMPthresh = 0.5) (Huppert et al., 2009). The optical-density signals were consequently band-pass filtered to include frequencies between 0.01 and 0.09 Hz in order to reduce systemic physiological artifacts (for example: cardiac, eye movement, respiratory and pulse artifacts). Signals were then converted from optical-density units to relative changes in hemoglobin concentration (HbO and HbR) using the modified Beer–Lambert law. Finally, we applied the block average approach (hmrBlockAvg function) to calculate the block average for HbO in each block and for each subject over the time window, which comprised—starting from a 2-s prestimulus—the stimulation duration and, in addition, 15 s to return to baseline. Then, the mean task-related changes of HbO referring to a 5-s baseline interval prior to the task (seconds -5 to 0) were calculated for each channel. The magnitude was extracted from the mean task-related change of the HbO curve during the stimulation duration and 5 s after its end.

### Statistical analyses

A 2×4 repeated measure analysis of variance (ANOVA) for repeated measurements (2 hemispheres [left vs right] × 4 stimulation conditions [8s, 10s, 15s and 20s]) was computed for the HbO parameter. If violation of the sphericity assumption occurred, Greenhouse–Geisser correction values were used for further analysis. To test the within-subject effect, we performed the paired sample t-tests. To account for multiple comparisons, the Bonferroni correction was employed.

In the second experiment, a t-test was used to determine the significant differences in cortical response to white noise and vocal/nonvocal sounds.

## Results

### Localization of cortical response to auditory stimulation: channel selections

In a preliminary step, given the binaural auditory stimulation conditions in our study, we estimated the MNI coordinates of the channels in our montage with the help of the “Project probe to cortex” tool available in AtlasViewer to ensure that the channels chosen for the study pass through the auditory cortex in the superior temporal gyrus. The resulting coordinates and their anatomical labels are provided in Table 1. Based on the table, one can conclude that the estimated anatomical regions, in which we measured the relative change in HbO concentration, correspond mostly to the superior temporal regions and the lateral sulcus, where the auditory areas are located (Penhune et al., 1996).

In the first step of analysis, which was based on data from all conditions of block stimulation durations (8s, 10s, 15s, and 20s) pooled together, we normalized all channels in the left and right hemispheres, respectively, to the most-activate channel. This normalization was performed separately on each side and the most-activate channel in each hemisphere taken as the maximum 100% value. There was a large range of activity across channels, as illustrated in Fig. 4. Given this significant variability across channels, we arbitrarily decided on a threshold of 65% to define the channels of interest to focus on as those with important activity and to filter out possible insignificant fluctuations. The reason for this particular threshold value was the retrieval of the same number of channels in each hemisphere. Therefore, all channels above 65% of the maximum response were selected and subjected to further analysis.

**Fig. 4.**
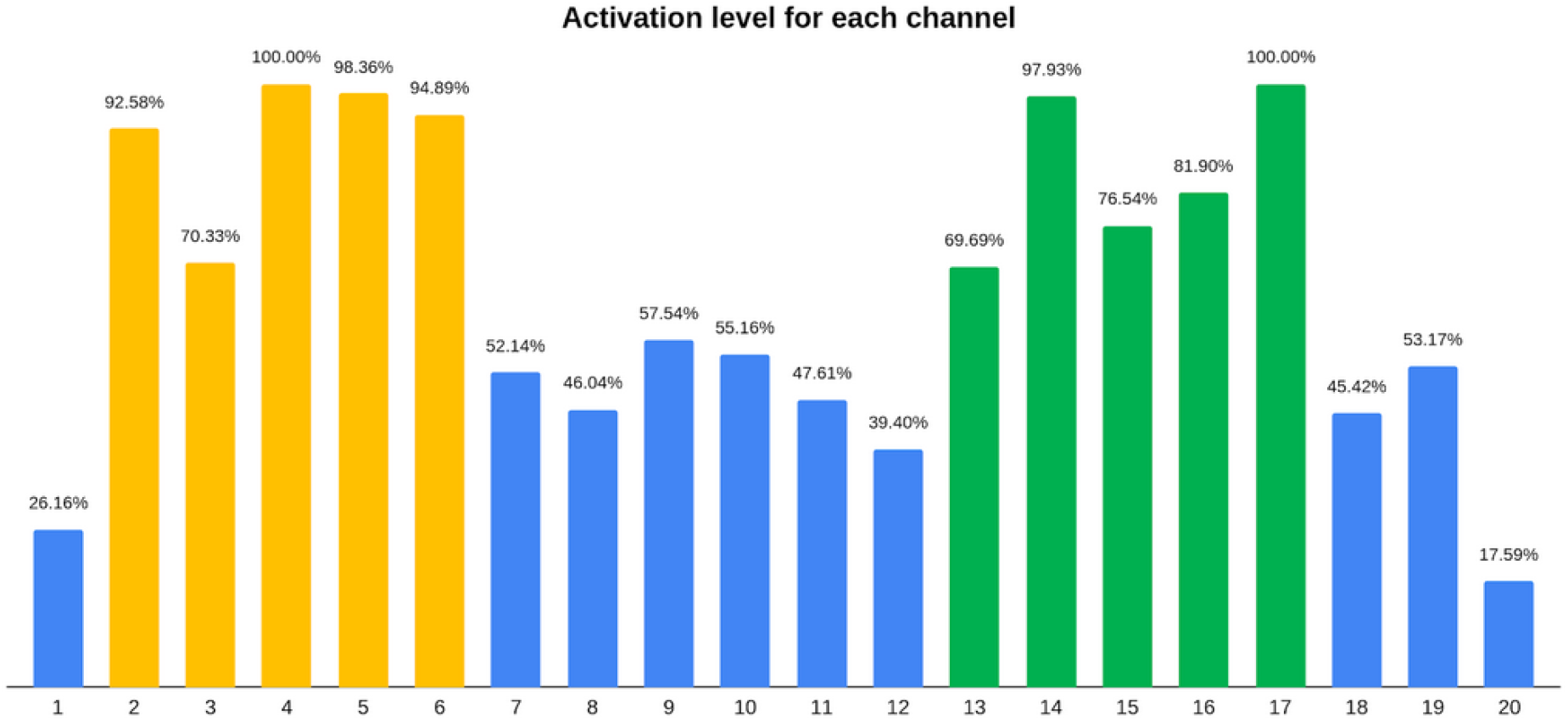
Activation level for each channel. The percentage of activation level with respect to the most active channel (100%) of each hemisphere. The x axis indicates the number of the channel. Channels from 1 to 10 are in the LH and channels from 11 to 20 are in the RH. The channels in yellow passed the activation threshold for the LH (ch:2-6) and in green for the RH (ch:13-17).

From the set of 10 channels in each hemisphere (Fig. 4), the most-activate channels chosen in this way were ch 2, 3, 4, 5, 6 in the left hemisphere (LH) and ch 13, 14, 15, 16, 17 in the right hemisphere (RH).

As expected due to the auditory nature of stimulation, the channels presenting the highest responses were localized in the auditory regions within the superior and middle temporal cortices, comprising the primary, secondary, and tertiary auditory areas (BA41, 21, and 22). It is worth noting that channel 4 had a 100% activation level and channel 5 had 98.36%, covering the superior temporal cortex, respectively (BA41, 21). Similarly, channels 14 and 17 had a 100% and 97.83% activation level, respectively, targeting the superior temporal cortex (BA41, 22) in the RH (Table 2). In addition, supramarginal (BA40) and precentral (BA6) regions were covered by the most-active channels.

**Table 2.** MNI coordinates and related Brodmann and anatomical areas of the channel-points.

| channel | src | det | MINI coord | Label name | BA |
| --- | --- | --- | --- | --- | --- |
| 1 | 1 | 1 | -54 12 8 | Rolandic_Oper_L | Left-BA44 |
| 2 | 1 | 2 | -60 5 23 | Precentral_L | Left-BA6 |
| 3 | 2 | 1 | -40 -6 -5 | Temporal_Sup_L | Left-Insula(13) |
| 4 | 2 | 2 | -48 -14 9 | Temporal_Sup_L | Left-PrimAuditory(41) |
| 5 | 2 | 3 | -49 -26 -6 | Temporal_Mid_L | Left-BA21 |
| 6 | 3 | 2 | -45 -22 26 | SupraMarginal_L | Left-Outside BAs |
| 7 | 3 | 3 | -60 -39 13 | Temporal_Mid_L | Left-BA22 |
| 8 | 3 | 4 | -51 -50 27 | Angular_L | Left-Outside BAs |
| 9 | 4 | 3 | -40 -45 -7 | Temporal_Inf_L | Left-Outside BAs |
| 10 | 4 | 4 | -57 -62 14 | Temporal_Mid_L | Left-BA39 |
| 11 | 5 | 5 | 59 12 8 | Frontal_Inf_oper_R | Right-BA44 |
| 12 | 5 | 6 | 56 1 -6 | Temporal_Sup_R | Right-BA22 |
| 13 | 6 | 5 | 56 7 22 | Frontal_Inf_oper_R | Right-BA44 |
| 14 | 6 | 6 | 60 -10 7 | Heschl_R | Right-PrimAuditory(41) |
| 15 | 6 | 7 | 60 -21 26 | SupraMarginal_R | Right-BA40 |
| 16 | 7 | 6 | 53 -26 -8 | Temporal_Mid_R | Right-BA21 |
| 17 | 7 | 7 | 65 -40 13 | Temporal_Sup_R | Right-BA22 |
| 18 | 8 | 7 | 60 -47 -2 | Temporal_Sup_R | Right-BA39 |
| 19 | 8 | 7 | 50 -48 25 | Angular_R | Right-AngGyrus(39) |
| 20 | 8 | 8 | 51 -58 16 | Temporal_id_R | Right-AngGyrus(39) |

Thus, the validity of the experimental technique, as well as the validity of the selection procedure for channels of interest, were confirmed by our results, the strongest responses to auditory stimulus being elicited in the auditory areas.

### Effects of auditory stimulation durations on the channels of interest

We estimated the influence of stimulation duration on the magnitude of the relative change in HbO concentration in each hemisphere with the 2*4 ANOVA for repeated measurements (hemispheres [left vs right] × 4 stimulation conditions [8s, 10s, 15s, and 20s]). This analysis demonstrated the highly significant main effect of the block stimulation duration (F[3; 132] = 10.881, p = 0.000146) on relative changes in HbO concentration with no differences between the hemispheres as determined by the insignificant hemispherical effect (F[1; 44] = 2.843, p>0.05) together with the insignificant interaction between stimulation conditions and hemispheres (F[3; 132] = 2.385, p>0.05).

Fig. 5 depicts how the amplitude of the averaged HDRs changes within the superior temporal regions and the lateral sulcus, detected by channels of interest with regard to stimulation durations. According to the post-hoc paired comparisons, the measured signal increased significantly from 8s to 10s and from 15s to 20s. The smaller increases in the signal (between 8s and 10s as well as between 15s and 20s) were not significantly different. For example and by way of illustration of the results in the bar plot, Fig. 6 shows the averaged signal time courses for 8s, 10s, 15s, and 20s conditions for the most-active channel, channel 15.

**Fig. 5.**
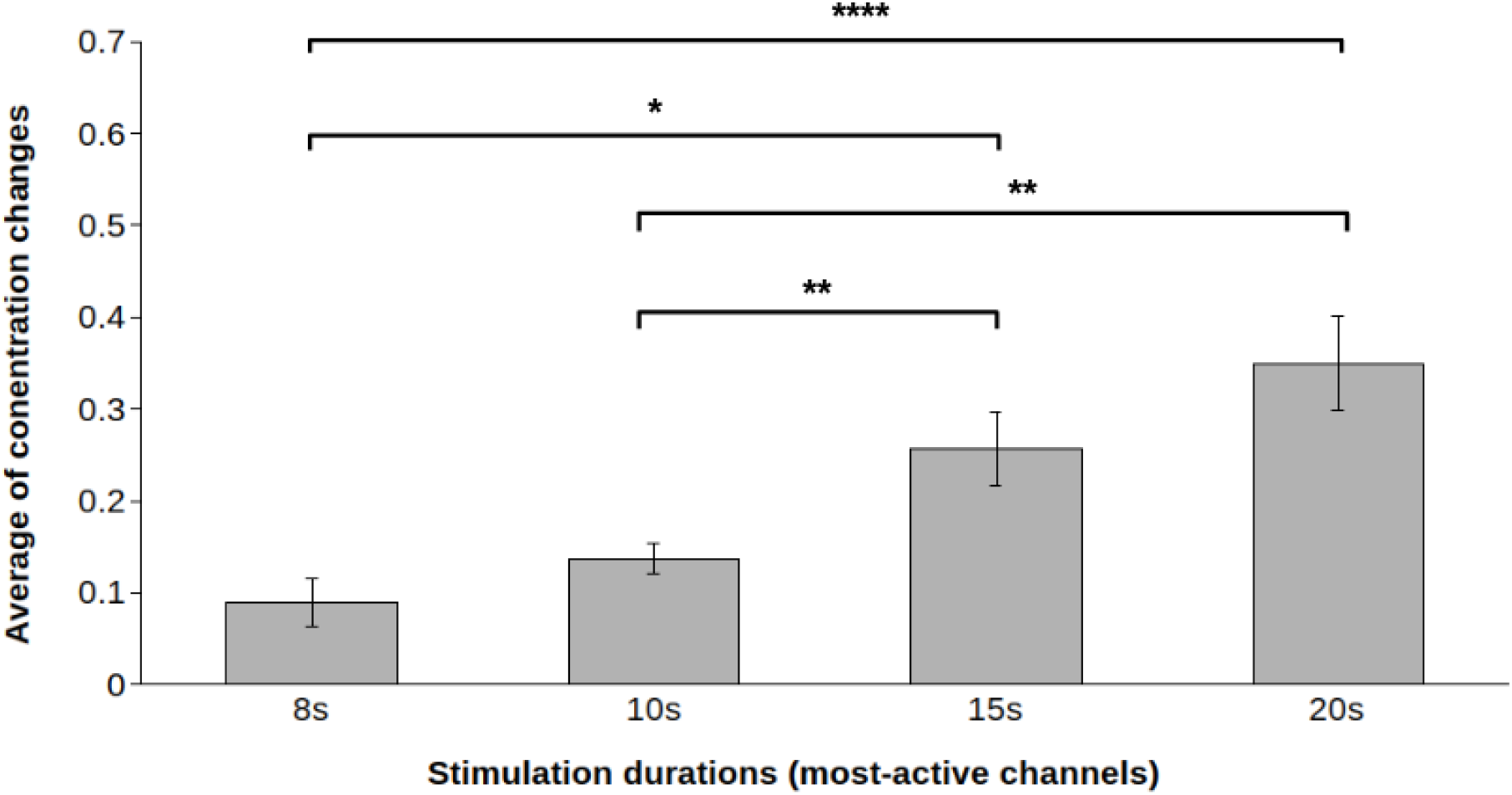
Effects of stimulation durations on the relative change in HbO concentration within the superior temporal regions and the lateral sulcus. The X axis represents four stimulation conditions and the Y axis indicates the amplitude of averaged changes of concentration on HbO taken from the most active channels detecting the auditory cortices. The significance of the ANOVA is indicated by stars: *p<0.05, **p<0.001, ****p<0.0001. The unit of measure on the Y axis is 10-7 mMol/L, and error bars represent standard errors.

**Fig. 6.**
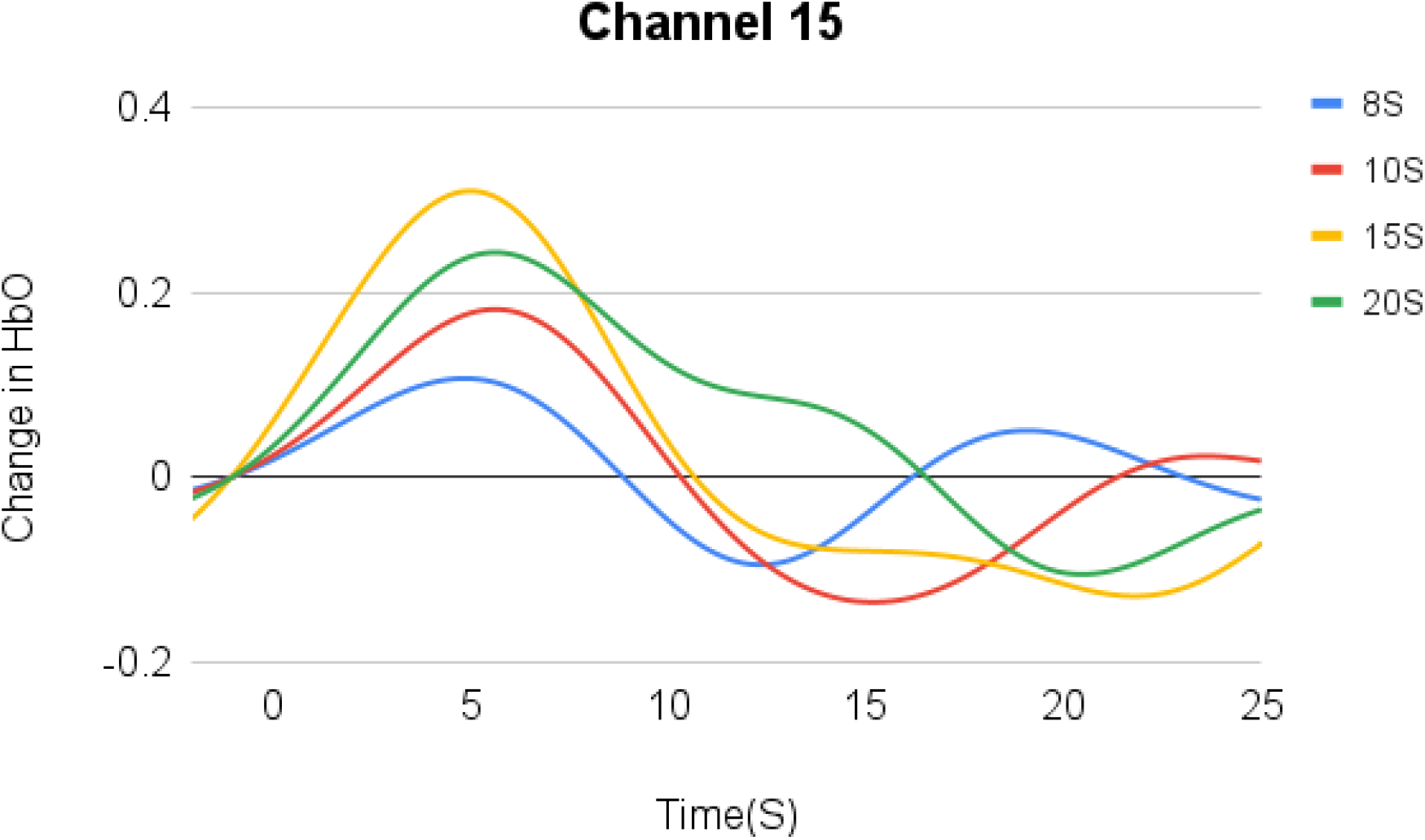
The averaged HDR time courses for four stimulation durations in channel 15. The X axis indicates the time courses in seconds for the 8s, 10s, 15s, and 20s stimulation conditions and the Y axis stands for the change in HbO concentration. The unit of measure on the Y axis is 10-7 mMol/L. The time courses to different durations are color encoded.

These results suggest that the hemodynamic response to auditory stimuli increases significantly with the increment of at least 5s of stimulation duration; however, the response appears to reach a plateau after 15s of stimulation. In general, as one can estimate visually in Fig. 5, the longer the stimulation duration, the greater the auditory cortical activation measured by fNIRS.

### Linearity analysis

A linear system model is defined by the principle of superposition. If the cortical activation level in response to a given simulation task f(x1+x2) can be predicted by the summation of the cortical activation level to individual tasks f(x1) and f(x2), this system follows a linear system model.

In our study, the additivity test was used to assess the linearity of the HDR with regard to the stimulation durations. To do so, we summed the responses to the shorter stimuli so as to predict the responses to the longer stimuli. Thus, the cortical response to 8s and 10s was doubled to estimate the response to 15s and 20s respectively. For example, the change in HbO with the duration of 10s was shifted by 10s and added to the 10s response to predict the change in HbO to the 20s stimulation. To assess the degree of goodness-of-fit between the predicted responses and the observed responses, the coefficient of determination was calculated using the complete time courses. The closer the value of this coefficient to one, the better the goodness-of-fit is and, therefore, the consistency with the assumption of linearity. Only the most-active channels in the primary auditory cortex were used for linearity analysis.

Fig. 7 depicts the predicted linear system responses overlaid with the observed responses for the left and right primary auditory cortices. The prediction for the 15s HDR time course based on the 8s HDR time course in the left and right auditory cortex exhibited differences with the measured response, as illustrated in Fig. 7a and 7b, confirmed by the low value of the coefficient of determination: R^2^ = 0.67 (left) and R^2^ = 0.17 (right). Afterward, the HDR time courses with a duration of 20s were predicted using a superposition of the HDR response to 10s in the left and right auditory cortices. As shown in Fig. 7c and 7d, the predicted responses are not well-fitted with the measured responses; thus, the conclusion is strengthened by the weak values of coefficients of determination: R^2^ = 0.38 (left) and R^2^ = 0.28 (right). The low value of this coefficient means that the superposition principle is not suitable for predicting the longer stimulation responses using the shorter stimulation signals (8s and 10s) for prediction; the system is nonlinear, at least for stimulation responses from 8s to 15s and from 10s to 20s.

**Fig. 7.**
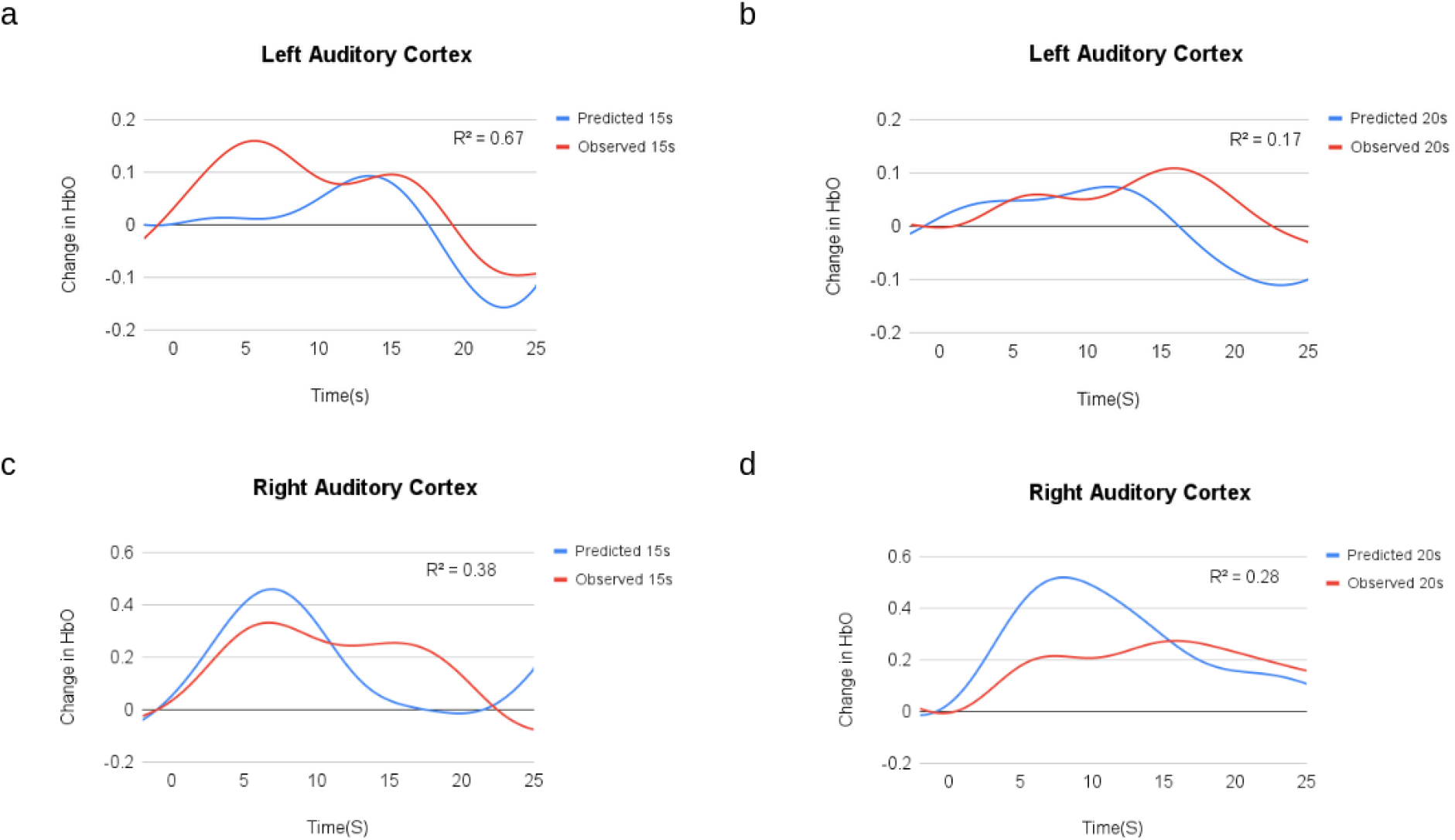
Predicted linear system responses in the auditory cortices compared with measured responses. (**a**) 15s linear system response established from superposition of two 8s responses, overlaid with observed response in the left auditory cortex. (**b**) 20s linear system response formed from superposition of two 20s responses, overlaid with measured responses in the left auditory cortex. (**c**) 15s linear system response overlaid with observed response in the right auditory cortex. (**d**) 20s linear system response overlaid with measured response in the right auditory cortex. The coefficient of determination(R^2^) is indicated in each plot, reflecting the degree of nonlinearity between the predicted response and observed response. Red lines indicate the measured hemodynamic responses and blue lines depict those predicted by the linear system responses.

### Effect of stimulus type: white noise vs natural sounds

Using the t-test analysis, we found no significant differences in the levels of cortical activation induced by white noise and those induced by natural sounds (non voices and voices) in our study (p>0.05). This suggests that auditory regions covered by the selected channels of interest respond to both types of stimuli in the same way (Fig. 8).

**Fig. 8.**
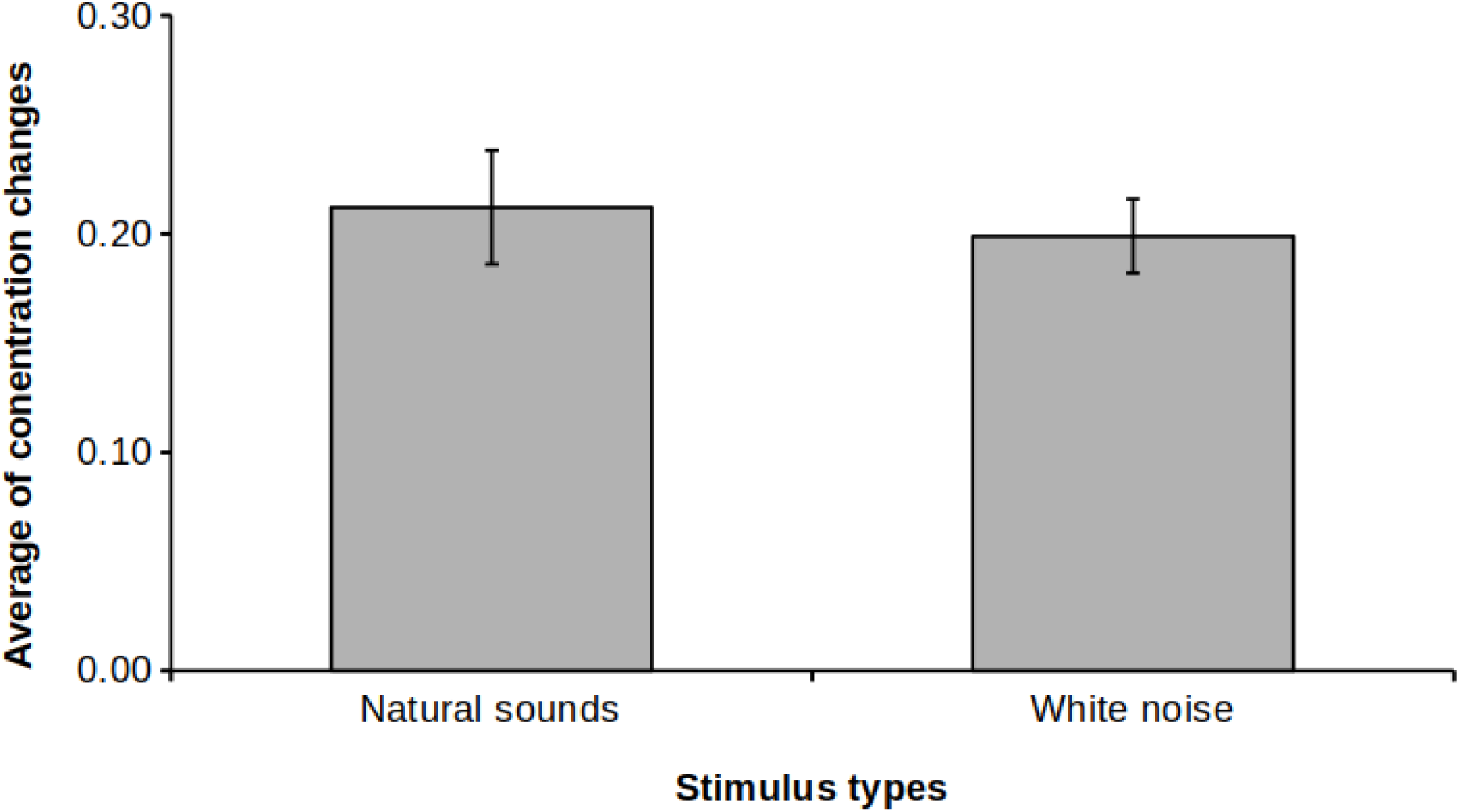
Effect of stimulus type on the HbO amplitude change in the superior temporal regions and the lateral sulcus. The horizontal axis represents two types of auditory stimulus: natural sounds and white noise. The average of the HbO concentration changes is displayed on the vertical axis. The unit of measure is 10^−7^ mMol/L, error bars represent standard deviations.

## Discussion

### Overview of results

The principal aim of this study was to evaluate the effects of different train stimulation durations on the auditory cortex hemodynamic response amplitude, thereby allowing the identification of the most-efficient stimulation periods for the auditory task using fNIRS. We observed that the auditory stimuli produce a well-localized cortical response in the superior temporal cortex, confirming the validity of our experimental design. The results further revealed that hemodynamic responses increase significantly with the increment of 5s of stimulation durations; however, the response reaches a plateau after 15s of stimulation. Afterward, we assessed the linearity of hemodynamic responses to different stimulation durations in the auditory cortices detected by the most-active channels. The linear superposition of responses of shorter duration failed to predict the response of longer duration, suggesting a nonlinear relationship for the range of our stimulation durations. The highest responses were observed in the conditions of 15s and 20s stimulation, and both were associated with a relatively short experiment duration. To avoid a saturated response at 20s, we chose a condition of 15s as the best stimulation duration to determine whether the peak amplitude is maintained with other types of more ecological stimuli, and the results showed that the stimulus type had no impact.

### Experimental validity: anatomic-functional correspondence

In our study, we demonstrated an anatomic-functional correspondence in the superior temporal cortex, validating fNIRs for auditory perception, in general, and more specifically, our experimental paradigm. In line with the literature findings using fMRI (e.g., (Hwang et al., 2007; Samson et al., 2011; Alho et al., 2014)), bilateral stimulation with trains of white noise induced the highest activations in the auditory cortices, and no significant differences were observed in the left and right hemispheres. In addition, a good sensibility of our optode setup to auditory cortical regions, confirmed by the results of Monte Carlo simulation for photon migration, as shown in Fig. 3, strengthens this anatomic-functional correspondence. These results validated the experimental setup for the following experiment.

### Stimulation durations and cortical response amplitude

Our findings also showed that the fNIRS response peak amplitudes to trains of white noise increased significantly with the increment of 5s in their duration and reached a plateau at 20s of stimulation duration. This observation suggests that hemodynamic responses in the auditory cortices become saturated for longer durations starting from about 15s.

Previous studies of the auditory response for different stimulation durations used a very large range of auditory stimulation durations, e.g., from 100ms to 25.5s (Robson et al., 1998), 1s to 16s (Soltysik et al., 2004), and 16s to 167s (Glover, 1999). The peak amplitudes for the auditory response increased with stimulation durations in all of those studies and reached a plateau at 8s in one study (Glover, 1999) and at 4s in another (Soltysik et al., 2004). Two more-recent studies (Hu et al., 2010; Ranaweera et al., 2016) on the effects of variable duration, both of which assessed the impact of acoustic noise on auditory hemodynamic responses, reported that, when the imaging acoustic noise duration increased, the induced hemodynamic responses also increased in amplitude in the auditory cortices. Hence, except for small variations in the timing where the responses reached a plateau, the cortical responses measured in our study are consistent with the findings of previous research using fMRI. The different experimental setups (such as the type of stimulus, the stimulus presentation rate, and sampling procedure of hemodynamic response) may explain the variations in response amplitude and duration across those studies.

### Hemodynamic response nonlinearity

The assessment of linearity in the left and right primary auditory cortices at the sites of the most-active channels showed the responses to trains of white noise to be mostly nonlinear, as evidenced by the weak value of the coefficient of determination between the linear system responses and measured responses. However, the degree of nonlinearity is higher for the longer stimulation duration of 20s than for 15s (modeled by the superposition of two responses to 10s and 8s respectively). It is worth noting that we applied the same quantitative measure of the linearity between the predicted response and the observed response and a similar model for the linearity assessment as that used in the previous studies using fMRI and fNIRS (Robson et al., 1998; Vazquez and Noll, 1998; Soltysik et al., 2004; Tian et al., 2009).

In general, our results confirm that the hemodynamic responses in the auditory cortex behave nonlinearly to trains of stimuli, especially for the longer duration. Previous studies indicated that the hemodynamic response exhibits strong nonlinearity for short stimulation durations, e.g., less than 7s (Soltysik et al., 2004) and less than 6s (Robson et al., 1998). Thus, hemodynamic responses to short-duration stimuli below this threshold cannot be used to predict the response to long stimulation. However, the stimuli of 8s and of 6s were approximate estimators to predict the behavior of longer stimuli for 16s (Soltysik et al., 2004), and 12.7s predicted 25.5s durations (Robson et al., 1998). Conversely, Glover indicated that the auditory response showed a somewhat more linear response for the short-duration stimuli but is highly nonlinear for longer stimuli (Glover, 1999). Taking these results into consideration, we suggest that auditory cortical responses to different stimuli durations are nonlinear in a global vision, but they can exhibit more or less linearity according to a certain range of stimulation durations.

Several investigators using fMRI attempted to model these nonlinearities using a Laplacian linear system to assess the BOLD response (Vazquez and Noll, 1998) and the neural adaptation functions, in which the influence of subsequent stimuli progressively decreased until it reached a steady level (Robson et al., 1998; Soltysik et al., 2004). Although these models have successfully predicted the nonlinear responses, they are not based on physiological parameters. However, the physiologically based balloon model and adaptive balloon model (Buxton et al., 1998, 2004; Havlicek et al., 2015) require estimating the contribution of various parameters, such as cerebral blood flow, cerebral blood volume, and cerebral metabolic oxygen rate, to the BOLD signal change. These contributing parameters are difficult to assess with current human neuroimaging methods.

Quantitative modeling of the hemodynamic response combined with experimental data facilitates the development of a quantitative assessment of brain physiology. Such modeling approaches have been applied extensively by fMRI studies and adapted in the present study. As fNIRS offers the possibility of the accurate and noninvasive measurement of the time course of HbO and HbR during human neuronal activation (Villringer et al., 1993), several studies (Huppert et al., 2006; Toyoda et al., 2008) combined the simultaneous measurement of fMRI and fNIRS. In this way, the authors tackled the quantitative evaluation of the contribution of oxygen extraction flow and cerebral blood volume to the BOLD signal using the model developed by Obata et al. (Obata et al., 2004) to elucidate brain physiology of these nonlinearities in the visual and motor cortex. Consequently, we believe that a good knowledge of nonlinearity in auditory fNIRS signals will provide the possibility of exploring neuronal activations based on hemodynamic response.

### Optimization aspects of stimulation parameters

Attempts to assess linearity or nonlinearity between the stimulation durations and the cortical response aim to optimize experimental design and improve the interpretation of brain activation. Given the described nonlinearity, our study demonstrated that the most-efficient stimulation duration is 15s with seven repeat trials for the presentation of about 100 stimuli. The conditions of 15s and 20s produced the highest amplitude of auditory cortical activation compared to other conditions. Although the amplitude at 20s is similar to the 15s condition and the former is associated with the shortest experimental duration, as shown in Table 1, the 15s condition may be a better choice. The nonlinearity analysis confirmed that the fNIRS signal exhibited higher nonlinearity and reached a plateau between 15s and 20s stimulation, suggesting increasing neuronal adaptation in this range of stimulation. To avoid a saturated response at 20s, we propose the 15s stimulation duration with seven repetitions as the most-suitable parameter of stimulation for block design in auditory studies using fNIRS.

### Effects of stimulus type

Having chosen the 15s stimulation duration as optimal, we compared white noise and natural sounds in a separate cohort. No significant differences were identified in terms of activation amplitude between white noise and natural sounds. White noise is considered relatively simple sound because of its variation only in the spectral dimension. Conversely, natural sounds are complex with regard to both their spectral and temporal variations (Samson et al., 2011). Both kinds of stimuli largely activate the superior temporal cortex and surrounding regions without interhemispheric lateralization, according to evidence from the meta-analysis of fMRI studies (Samson et al., 2011). Compared to white noise, natural sounds, including vocal and nonvocal sounds, exhibit more-ecological characteristics. These results confirmed the validity of the choice of the 15s duration for more-complex stimuli. Most importantly, our study demonstrated that the strongest activation in the 15s condition is not dependent on stimulus type, thereby reinforcing the reliability of our conclusions.

### Limitations of the method

Compared to fMRI, the spatial resolution of fNIRS is low. The trajectory of the light beam from emitter to detector is thought to cover the underlying cortical region (Okada et al., 1997), but how deeply into the brain tissue the fNIRS measure spreads is unclear (Cui et al., 2011). Consequently, the primary auditory cortex may not be completely detectable due to the limited penetration depth. In addition, the sampling volume in fNIRS corresponds to multiple voxels in fMRI, leading to cruder measurements in terms of spatial resolution.

Another point worth mentioning involves the differences in hemodynamic time courses in Fig. 7. The temporal variability of the hemodynamic response (i.e., in peaks, shape, and amplitude) is a frequent BOLD phenomenon in fMRI studies (Buckner, 1998; Buckner et al., 1998). Several reasons for this variability are identified in the literature, such as a delay in the underlying neural implication, vasculature differences, physiological differences, level of consciousness, etc.(Jasdzewski et al., 2003; Handwerker et al., 2004; Erdoğan et al., 2016; Kamran et al., 2016).

### Practical applications in fNIRs studies

In practice, increasing the number of trials and the intertrial intervals leads to significant increases in the experimental time, while decreasing the intertrial intervals decreases the amplitude induced by the subsequent stimulation trial (hemodynamic refractory effect) (Inan et al., 2004). Additionally, decreasing the number of trials decreases the signal-to-noise ratio, suggesting that a conflict exists between the experimental time and the quality of the signal. How can stimulation be conducted most efficiently with a relatively short experimental duration and the maximal cortical activation? Is there a most-optimal organization of distribution of a given number of stimuli? Our results demonstrate that a combination of relatively longer stimulation duration with a relatively smaller number of trials is more efficient for stimulating the auditory cortex than the combination of a relatively shorter stimulation duration with a larger number of trials, suggesting that the effects of stimulation duration on fNIRS signal quality are more pronounced than the number of trials. This is a critical point to consider for experimental block design in fNIRS studies. It is worth mentioning that, as the dominant application of fNIRS research involves infants and children, as well as neuropsychiatric populations (Boas et al., 2014), investigators should consider longer stimulation durations and fewer trials in their studies.

### Perspectives for future research

Given the practical advantages of fNIRS, such as ease of use and low cost, more fine-tuned assessments of stimulation durations with short incremental intervals should be tested in the future to refine the range of linear and nonlinear responses to stimuli durations. The presence of nonlinearity in the hemodynamic response suggests that multimodal experiments should be conducted in order to better understand the nature of the relationship between neural activity and the measured signal. The simultaneous measurement of fMRI, EEG, and fNIRS will make it possible to investigate in greater detail the neural origins of nonlinearity in the auditory cortical responses to different stimulation durations. In addition to temporal aspects of brain activity, spatial patterns of auditory cortical responses to different stimulation durations can be investigated (Sadoun et al., 2020).

## Conclusions

The results of the current study indicate that the fNIRS signal is sensitive to differences in stimulation duration, emphasizing the importance of considering block duration effects in auditory fNIRS experimental designs. We demonstrated that hemodynamic responses increase significantly due to stimulation duration increments and reach a plateau at the block duration of 15s, with the highest values for the durations of 15s and 20s. The linearity analysis showed that the increase in activity for the longer stimulation duration is not linear in the auditory cortex. To avoid a saturated response at 20s, we chose the 15s condition as the most-efficient stimulation duration, independent of stimulation type. Since NIRS has become an increasingly popular method in the field of hearing research, the optimization of experimental design that we aimed at in our study should help establish more-efficient experimental designs for the high quality of the fNIRS signals.

## Abbreviations

deoxy-Hb: deoxyhemoglobin
fNIRS: functional near-infrared spectroscopy
fMRI: functional magnetic resonance imaging
HbO: Oxygenated haemoglobin
HbR: Deoxygenated hemoglobin
HbT: Total hemoglobin concentration
HDR: hemodynamic response
LH: Left hemisphere
oxy-Hb: oxyhemoglobin
RH: Right hemisphere
SNR: signal-to-noise ratio

## Acknowledgements

We would like to thank Dr. Kuzma Strelnikov for helpful advice regarding data analysis.

## Authors’ contributions

YFZ : designed the experiments, performed all the data analysis and statistical analyses, also the manipulations and wrote the manuscript. AL : technical support. IB: project management. All authors read and approved the final manuscript.

## References

Aasted, C.M., Yücel, M.A., Cooper, R.J., Dubb, J., Tsuzuki, D., Becerra, L., Petkov, M.P., Borsook, D., Dan, I., Boas, D.A., 2015. Anatomical guidance for functional near-infrared spectroscopy: AtlasViewer tutorial. Neurophotonics 2, 020801. https://doi.org/10.1117/1.NPh.2.2.020801

Alho, K., Rinne, T., Herron, T.J., Woods, D.L., 2014. Stimulus-dependent activations and attention-related modulations in the auditory cortex: a meta-analysis of fMRI studies. Hear Res 307, 29–41. https://doi.org/10.1016/j.heares.2013.08.001

Amaro, E., Barker, G.J., 2006. Study design in fMRI: Basic principles. Brain and Cognition 60, 220–232. https://doi.org/10.1016/j.bandc.2005.11.009

Anderson, C.A., Wiggins, I.M., Kitterick, P.T., Hartley, D.E.H., 2019. Pre-operative Brain Imaging Using Functional Near-Infrared Spectroscopy Helps Predict Cochlear Implant Outcome in Deaf Adults. J Assoc Res Otolaryngol 20, 511–528. https://doi.org/10.1007/s10162-019-00729-z

Basura, G.J., Hu, X.-S., Juan, J.S., Tessier, A.-M., Kovelman, I., 2018. Human central auditory plasticity: A review of functional near-infrared spectroscopy (fNIRS) to measure cochlear implant performance and tinnitus perception. Laryngoscope Investig Otolaryngol 3, 463–472. https://doi.org/10.1002/lio2.185

Bauernfeind, G., Wriessnegger, S.C., Haumann, S., Lenarz, T., 2018. Cortical activation patterns to spatially presented pure tone stimuli with different intensities measured by functional near-infrared spectroscopy. Human Brain Mapping 39, 2710–2724. https://doi.org/10.1002/hbm.24034

Belin, P., Zatorre, R.J., Lafaille, P., Ahad, P., Pike, B., 2000. Voice-selective areas in human auditory cortex. Nature 403, 309–312. https://doi.org/10.1038/35002078

Binder, J.R., Rao, S.M., Hammeke, T.A., Frost, J.A., Bandettini, P.A., Hyde, J.S., 1994. Effects of stimulus rate on signal response during functional magnetic resonance imaging of auditory cortex. Cognitive Brain Research 2, 31–38. https://doi.org/10.1016/0926-6410(94)90018-3

Boas, D., Culver, J., Stott, J., Dunn, A., 2002. Three dimensional Monte Carlo code for photon migration through complex heterogeneous media including the adult human head. Opt Express 10, 159–170. https://doi.org/10.1364/oe.10.000159

Boas, D.A., Dale, A.M., Franceschini, M.A., 2004. Diffuse optical imaging of brain activation: approaches to optimizing image sensitivity, resolution, and accuracy. Neuroimage 23 Suppl 1, S275–288. https://doi.org/10.1016/j.neuroimage.2004.07.011

Boas, D.A., Elwell, C.E., Ferrari, M., Taga, G., 2014. Twenty years of functional near-infrared spectroscopy: introduction for the special issue. Neuroimage 85 Pt 1, 1–5. https://doi.org/10.1016/j.neuroimage.2013.11.033

Boynton, G.M., Engel, S.A., Glover, G.H., Heeger, D.J., 1996. Linear Systems Analysis of Functional Magnetic Resonance Imaging in Human V1. J. Neurosci. 16, 4207–4221. https://doi.org/10.1523/JNEUROSCI.16-13-04207.1996

Buckner, R.L., 1998. Event-related fMRI and the hemodynamic response. Human Brain Mapping 6, 373–377. https://doi.org/10.1002/(SICI)1097-0193(1998)6:5/6<373::AID-HBM8>3.0.CO;2-P

Buckner, R.L., Koutstaal, W., Schacter, D.L., Wagner, A.D., Rosen, B.R., 1998. Functional-anatomic study of episodic retrieval using fMRI. I. Retrieval effort versus retrieval success. Neuroimage 7, 151–162. https://doi.org/10.1006/nimg.1998.0327

Buxton, R.B., Uludağ, K., Dubowitz, D.J., Liu, T.T., 2004. Modeling the hemodynamic response to brain activation. Neuroimage 23 Suppl 1, S220–233. https://doi.org/10.1016/j.neuroimage.2004.07.013

Buxton, R.B., Wong, E.C., Frank, L.R., 1998. Dynamics of blood flow and oxygenation changes during brain activation: the balloon model. Magn Reson Med 39, 855–864. https://doi.org/10.1002/mrm.1910390602

Cui, X., Bray, S., Bryant, D.M., Glover, G.H., Reiss, A.L., 2011. A quantitative comparison of NIRS and fMRI across multiple cognitive tasks. Neuroimage 54, 2808–2821. https://doi.org/10.1016/j.neuroimage.2010.10.069

Erdoğan, S.B., Tong, Y., Hocke, L.M., Lindsey, K.P., deB Frederick, B., 2016. Correcting for Blood Arrival Time in Global Mean Regression Enhances Functional Connectivity Analysis of Resting State fMRI-BOLD Signals. Front Hum Neurosci 10, 311. https://doi.org/10.3389/fnhum.2016.00311

Fava, E., Hull, R., Baumbauer, K., Bortfeld, H., 2014. Hemodynamic responses to speech and music in preverbal infants. Child Neuropsychol 20, 430–448. https://doi.org/10.1080/09297049.2013.803524

Ferrari, M., Quaresima, V., 2012. A brief review on the history of human functional near-infrared spectroscopy (fNIRS) development and fields of application. NeuroImage 63, 921–935. https://doi.org/10.1016/j.neuroimage.2012.03.049

Gagnon, L., Yücel, M.A., Dehaes, M., Cooper, R.J., Perdue, K.L., Selb, J., Huppert, T.J., Hoge, R.D., Boas, D.A., 2012. Quantification of the cortical contribution to the NIRS signal over the motor cortex using concurrent NIRS-fMRI measurements. Neuroimage 59, 3933–3940. https://doi.org/10.1016/j.neuroimage.2011.10.054

Glover, G.H., 1999. Deconvolution of impulse response in event-related BOLD fMRI. Neuroimage 9, 416–429. https://doi.org/10.1006/nimg.1998.0419

Handwerker, D.A., Ollinger, J.M., D’Esposito, M., 2004. Variation of BOLD hemodynamic responses across subjects and brain regions and their effects on statistical analyses. Neuroimage 21, 1639–1651. https://doi.org/10.1016/j.neuroimage.2003.11.029

Havlicek, M., Roebroeck, A., Friston, K., Gardumi, A., Ivanov, D., Uludag, K., 2015. Physiologically informed dynamic causal modeling of fMRI data. Neuroimage 122, 355–372. https://doi.org/10.1016/j.neuroimage.2015.07.078

Herold, F., Wiegel, P., Scholkmann, F., Müller, N.G., 2018. Applications of Functional Near-Infrared Spectroscopy (fNIRS) Neuroimaging in Exercise–Cognition Science: A Systematic, Methodology-Focused Review. J Clin Med 7. https://doi.org/10.3390/jcm7120466

Hiwa, S., Katayama, T., Hiroyasu, T., 2018. Functional near-infrared spectroscopy study of the neural correlates between auditory environments and intellectual work performance. Brain Behav 8, e01104. https://doi.org/10.1002/brb3.1104

Homan, R.W., Herman, J., Purdy, P., 1987. Cerebral location of international 10-20 system electrode placement. Electroencephalogr Clin Neurophysiol 66, 376–382. https://doi.org/10.1016/0013-4694(87)90206-9

Hu, S., Olulade, O., Gonzalez, J.C., Santos, J., Kim, S., Tamer, G.G., Luh, W.-M., Talavage, T.M., 2010. Modeling Hemodynamic Responses in Auditory Cortex at 1.5T Using Variable Duration Imaging Acoustic Noise. Neuroimage 49, 3027. https://doi.org/10.1016/j.neuroimage.2009.11.051

Huppert, T.J., Diamond, S.G., Franceschini, M.A., Boas, D.A., 2009. HomER: a review of time-series analysis methods for near-infrared spectroscopy of the brain. Appl Opt 48, D280–298. https://doi.org/10.1364/ao.48.00d280

Huppert, T.J., Hoge, R.D., Diamond, S.G., Franceschini, M.A., Boas, D.A., 2006. A temporal comparison of BOLD, ASL, and NIRS hemodynamic responses to motor stimuli in adult humans. Neuroimage 29, 368–382. https://doi.org/10.1016/j.neuroimage.2005.08.065

Hwang, J.-H., Li, C.-W., Wu, C.-W., Chen, J.-H., Liu, T.-C., 2007. Aging effects on the activation of the auditory cortex during binaural speech listening in white noise: an fMRI study. Audiol Neurootol 12, 285–294. https://doi.org/10.1159/000103209

Inan, S., Mitchell, T., Song, A., Bizzell, J., Belger, A., 2004. Hemodynamic correlates of stimulus repetition in the visual and auditory cortices: an fMRI study. Neuroimage 21, 886–893. https://doi.org/10.1016/j.neuroimage.2003.10.029

Issa, M., Bisconti, S., Kovelman, I., Kileny, P., Basura, G.J., 2016. Human Auditory and Adjacent Nonauditory Cerebral Cortices Are Hypermetabolic in Tinnitus as Measured by Functional Near-Infrared Spectroscopy (fNIRS). Neural Plast 2016, 7453149. https://doi.org/10.1155/2016/7453149

Issard, C., Gervain, J., 2018. Variability of the hemodynamic response in infants: Influence of experimental design and stimulus complexity. Dev Cogn Neurosci 33, 182–193. https://doi.org/10.1016/j.dcn.2018.01.009

Jasdzewski, G., Strangman, G., Wagner, J., Kwong, K.K., Poldrack, R.A., Boas, D.A., 2003. Differences in the hemodynamic response to event-related motor and visual paradigms as measured by near-infrared spectroscopy. Neuroimage 20, 479–488. https://doi.org/10.1016/s1053-8119(03)00311-2

Kamran, M.A., Mannan, M.M.N., Jeong, M.Y., 2016. Cortical Signal Analysis and Advances in Functional Near-Infrared Spectroscopy Signal: A Review. Front. Hum. Neurosci. 10. https://doi.org/10.3389/fnhum.2016.00261

Krekelberg, B., Boynton, G.M., van Wezel, R.J.A., 2006. Adaptation: from single cells to BOLD signals. Trends Neurosci 29, 250–256. https://doi.org/10.1016/j.tins.2006.02.008

Liu, H., Gao, J., 2000. An investigation of the impulse functions for the nonlinear BOLD response in functional MRI. Magn Reson Imaging 18, 931–938. https://doi.org/10.1016/s0730-725x(00)00214-9

Malonek, D., Grinvald, A., 1996. Interactions between electrical activity and cortical microcirculation revealed by imaging spectroscopy: implications for functional brain mapping. Science 272, 551–554. https://doi.org/10.1126/science.272.5261.551

Massida, Z., Belin, P., James, C., Rouger, J., Fraysse, B., Barone, P., Deguine, O., 2011. Voice discrimination in cochlear-implanted deaf subjects. Hear Res 275, 120–129. https://doi.org/10.1016/j.heares.2010.12.010

Miezin, F.M., Maccotta, L., Ollinger, J.M., Petersen, S.E., Buckner, R.L., 2000. Characterizing the Hemodynamic Response: Effects of Presentation Rate, Sampling Procedure, and the Possibility of Ordering Brain Activity Based on Relative Timing. NeuroImage 11, 735–759. https://doi.org/10.1006/nimg.2000.0568

Nangini, C., Staines, W.R., McIlroy, W.I., Graham, S., 2002. Non-linear Event-Related fMRI BOLD Responses in Human Somatosensory Cortex. Proc. Intl. Soc. Mag. Reson. Med. 1.

Obata, T., Liu, T.T., Miller, K.L., Luh, W.M., Wong, E.C., Frank, L.R., Buxton, R.B., 2004. Discrepancies between BOLD and flow dynamics in primary and supplementary motor areas: application of the balloon model to the interpretation of BOLD transients. Neuroimage 21, 144–153. https://doi.org/10.1016/j.neuroimage.2003.08.040

Okada, E., Firbank, M., Schweiger, M., Arridge, S.R., Cope, M., Delpy, D.T., 1997. Theoretical and experimental investigation of near-infrared light propagation in a model of the adult head. Appl Opt 36, 21–31. https://doi.org/10.1364/ao.36.000021

Peelle, J.E., 2014. Methodological challenges and solutions in auditory functional magnetic resonance imaging. Front Neurosci 8. https://doi.org/10.3389/fnins.2014.00253

Penhune, V.B., Zatorre, R.J., MacDonald, J.D., Evans, A.C., 1996. Interhemispheric anatomical differences in human primary auditory cortex: probabilistic mapping and volume measurement from magnetic resonance scans. Cereb Cortex 6, 661–672. https://doi.org/10.1093/cercor/6.5.661

Pfeuffer, J., McCullough, J.C., Van de Moortele, P.F., Ugurbil, K., Hu, X., 2003. Spatial dependence of the nonlinear BOLD response at short stimulus duration. Neuroimage 18, 990–1000. https://doi.org/10.1016/s1053-8119(03)00035-1

Pinti, P., Scholkmann, F., Hamilton, A., Burgess, P., Tachtsidis, I., 2019. Current Status and Issues Regarding Pre-processing of fNIRS Neuroimaging Data: An Investigation of Diverse Signal Filtering Methods Within a General Linear Model Framework. Front. Hum. Neurosci. 12. https://doi.org/10.3389/fnhum.2018.00505

Pollonini, L., Olds, C., Abaya, H., Bortfeld, H., Beauchamp, M.S., Oghalai, J.S., 2014. Auditory cortex activation to natural speech and simulated cochlear implant speech measured with functional near-infrared spectroscopy. Hear Res 309, 84–93. https://doi.org/10.1016/j.heares.2013.11.007

Ranaweera, R.D., Kwon, M., Hu, S., Tamer, G.G., Luh, W.-M., Talavage, T.M., 2016. Temporal pattern of acoustic imaging noise asymmetrically modulates activation in the auditory cortex. Hear Res 331, 57–68. https://doi.org/10.1016/j.heares.2015.09.017

Remijn, G.B., Kikuchi, M., Yoshimura, Y., Shitamichi, K., Ueno, S., Tsubokawa, T., Kojima, H., Higashida, H., Minabe, Y., 2017. A Near-Infrared Spectroscopy Study on Cortical Hemodynamic Responses to Normal and Whispered Speech in 3-to 7-Year-Old Children. J Speech Lang Hear Res 60, 465–470. https://doi.org/10.1044/2016_JSLHR-H-15-0435

Robson, M.D., Dorosz, J.L., Gore, J.C., 1998. Measurements of the Temporal fMRI Response of the Human Auditory Cortex to Trains of Tones. NeuroImage 7, 185–198. https://doi.org/10.1006/nimg.1998.0322

Sadoun, A., Chauhan, T., Mameri, S., Zhang, Y.F., Barone, P., Deguine, O., Strelnikov, K., 2020. Stimulus-specific information is represented as local activity patterns across the brain. Neuroimage 223, 117326. https://doi.org/10.1016/j.neuroimage.2020.117326

Samson, F., Zeffiro, T.A., Toussaint, A., Belin, P., 2011. Stimulus Complexity and Categorical Effects in Human Auditory Cortex: An Activation Likelihood Estimation Meta-Analysis. Front. Psychol. 1. https://doi.org/10.3389/fpsyg.2010.00241

Shan, Z.Y., Wright, M.J., Thompson, P.M., McMahon, K.L., Blokland, G.G.A.M., de Zubicaray, G.I., Martin, N.G., Vinkhuyzen, A.A.E., Reutens, D.C., 2014. Modeling of the hemodynamic responses in block design fMRI studies. J Cereb Blood Flow Metab 34, 316–324. https://doi.org/10.1038/jcbfm.2013.200

Soltysik, D.A., Peck, K.K., White, K.D., Crosson, B., Briggs, R.W., 2004. Comparison of hemodynamic response nonlinearity across primary cortical areas. NeuroImage 22, 1117–1127. https://doi.org/10.1016/j.neuroimage.2004.03.024

Steinbrink, J., Villringer, A., Kempf, F., Haux, D., Boden, S., Obrig, H., 2006. Illuminating the BOLD signal: combined fMRI-fNIRS studies. Magn Reson Imaging 24, 495–505. https://doi.org/10.1016/j.mri.2005.12.034

Strangman, G., Culver, J.P., Thompson, J.H., Boas, D.A., 2002. A quantitative comparison of simultaneous BOLD fMRI and NIRS recordings during functional brain activation. Neuroimage 17, 719–731.

Tian, F., Chance, B., Liu, H., 2009. Investigation of the prefrontal cortex in response to duration-variable anagram tasks using functional near-infrared spectroscopy. J Biomed Opt 14, 054016. https://doi.org/10.1117/1.3241984

Toyoda, H., Kashikura, K., Okada, T., Nakashita, S., Honda, M., Yonekura, Y., Kawaguchi, H., Maki, A., Sadato, N., 2008. Source of nonlinearity of the BOLD response revealed by simultaneous fMRI and NIRS. Neuroimage 39, 997–1013. https://doi.org/10.1016/j.neuroimage.2007.09.053

Vannson, N., Strelnikov, K., James, C.J., Deguine, O., Barone, P., Marx, M., 2020. Evidence of a functional reorganization in the auditory dorsal stream following unilateral hearing loss. Neuropsychologia 149, 107683. https://doi.org/10.1016/j.neuropsychologia.2020.107683

Vazquez, A.L., Noll, D.C., 1998. Nonlinear aspects of the BOLD response in functional MRI. Neuroimage 7, 108–118. https://doi.org/10.1006/nimg.1997.0316

Villringer, A., Planck, J., Hock, C., Schleinkofer, L., Dirnagl, U., 1993. Near infrared spectroscopy (NIRS): a new tool to study hemodynamic changes during activation of brain function in human adults. Neurosci Lett 154, 101–104. https://doi.org/10.1016/0304-3940(93)90181-j

Virtanen, J., Noponen, T., Meriläinen, P., 2009. Comparison of principal and independent component analysis in removing extracerebral interference from near-infrared spectroscopy signals. J Biomed Opt 14, 054032. https://doi.org/10.1117/1.3253323

Weder, S., Shoushtarian, M., Olivares, V., Zhou, X., Innes-Brown, H., McKay, C., 2020. Cortical fNIRS Responses Can Be Better Explained by Loudness Percept than Sound Intensity. Ear and Hearing 41, 1187–1195. https://doi.org/10.1097/AUD.0000000000000836

Weder, S., Zhou, X., Shoushtarian, M., Innes-Brown, H., McKay, C., 2018. Cortical Processing Related to Intensity of a Modulated Noise Stimulus—a Functional Near-Infrared Study. JARO 19, 273–286. https://doi.org/10.1007/s10162-018-0661-0

Wilcox, J.B., Howell, R.D., Breivik, E., 2008. Questions about formative measurement. Journal of Business Research, Formative Indicators 61, 1219–1228. https://doi.org/10.1016/j.jbusres.2008.01.010

Wilcox, T., Haslup, J.A., Boas, D.A., 2010. Dissociation of processing of featural and spatiotemporal information in the infant cortex. NeuroImage 53, 1256–1263. https://doi.org/10.1016/j.neuroimage.2010.06.064

Wobst, P., Wenzel, R., Kohl, M., Obrig, H., Villringer, A., 2001. Linear Aspects of Changes in Deoxygenated Hemoglobin Concentration and Cytochrome Oxidase Oxidation during Brain Activation. NeuroImage 13, 520–530. https://doi.org/10.1006/nimg.2000.0706

Ye, J.C., Tak, S., Jang, K.E., Jung, J., Jang, J., 2009. NIRS-SPM: statistical parametric mapping for near-infrared spectroscopy. Neuroimage 44, 428–447. https://doi.org/10.1016/j.neuroimage.2008.08.036

Zhang, M., Mary Ying, Y.-L., Ihlefeld, A., 2018. Spatial Release From Informational Masking: Evidence From Functional Near Infrared Spectroscopy. Trends Hear 22, 2331216518817464. https://doi.org/10.1177/2331216518817464

Zhang, Y., Brooks, D.H., Franceschini, M.A., Boas, D.A., 2005. Eigenvector-based spatial filtering for reduction of physiological interference in diffuse optical imaging. J Biomed Opt 10, 11014. https://doi.org/10.1117/1.1852552

